# annotatr: Genomic regions in context

**DOI:** 10.1101/039685

**Authors:** Raymond G. Cavalcante, Maureen A. Sartor

## Abstract

**Motivation:** Analysis of next-generation sequencing data often results in a list of genomic regions. These may include differentially methylated CpGs/regions, transcription factor binding sites, interacting chromatin regions, or GWAS-associated SNPs, among others. A common analysis step is to annotate such genomic regions to genomic annotations (promoters, exons, enhancers, etc.). Existing tools are limited by a lack of annotation sources and flexible options, the time it takes to annotate regions, an artificial one-to-one region-to-annotation mapping, a lack of visualization options to easily summarize data, or some combination thereof.

**Results:** We developed the annotatr Bioconductor package to flexibly and quickly summarize and plot annotations of genomic regions. The annotatr package reports all intersections of regions and annotations, giving a better understanding of the genomic context of the regions. A variety of graphics functions are implemented to easily plot numerical or categorical data associated with the regions across the annotations, and across annotation intersections, providing insight into how characteristics of the regions differ across the annotations. We demonstrate that annotatr is up to 27x faster than comparable R packages. Overall, annotatr enables a richer biological interpretation of experiments.

**Availability:** http://bioconductor.org/packages/annotatr/

**Supplementary information:** Supplementary data are available at *Bioinformatics* online.

## 1 Introduction

Genomic regions resulting from next-generation sequencing experiments and bioinformatics pipelines require annotation to genomic features for context. For example, hyper-methylation of CpG shores in promoters may indicate different regulatory regimes in one condition compared to another, or it may be of interest that a transcription factor overwhelmingly binds in enhancers, while another tends to bind at exon-intron boundaries.

While tools exist to intersect genomic regions of interest with genomic annotations, we found the annotations, methods of intersection, and graphics options had room for improvement. ChIPpeakAnno (Zhu *et al.*, 2010) is an R package that has been used in many studies across a variety of organisms. It returns only one genomic annotation per input region, and while providing the user with some plots, these are limited by their inability to incorporate data associated with the regions of interest such as methylation rates, fold changes, etc. Another R package, goldmine (Bhasin & Ting, 2016), returns either all annotations intersecting regions of interest (a one-to-many mapping) or annotations based on a prioritization (a one-to-one mapping). The goldmine package provides helper functions to create annotations from any UCSC genome browser table. However, it does not offer built-in functions for summary plots, nor to plot data related to the regions over the annotations. Outside of the R ecosystem, BEDtools (Quinlan & Hall, 2010), implemented in C++, intersects and aggregates genomic regions with annotations, and is very fast. However, its more general purpose means users must provide all annotations and manually generate plots.

We developed annotatr, a Bioconductor package that reports all intersections of genomic regions with built-in genomic annotations for *D. melanogaster* (dm3 and dm6), *H. Sapiens* (hg19 and hg38), *M. musculus* (mm9 and mm10), *R. norvegicus* (rn4, rn5, and rn6), or custom annotations for any organism. annotatr enables users to associate numerical or categorical data with regions, enabling better understanding of the underlying experiments via summarization and visualization functions. annotatr is fast, flexible, and easily included in bioinformatics pipelines.

## 2 Implementation and Features

A core feature of annotatr is the variety of standard and specialized genomic annotations it includes. Standard annotations include CpG island related features (CpG islands, shores, shelves, and “open sea”) and genic features (promoters, 5’UTRs, exons, introns, CDS, and 3’UTRs) (Figure S1). Specialized genomic annotations include intron/exon boundaries, enhancers, lncRNAs, and chromatin state segmentations. Details regarding the sources, construction, and genome availability for all included annotations are provided in Supplementary Methods and Table S1. Custom annotations can supplement supported organisms, or enable annotation to any organism not listed above.

The annotatr package consists of four modules that read, annotate, summarize, and visualize genomic regions. The read module reads a BED6+ file, defined as BED6 and any number of numerical or categorical data columns. The annotate module reports the overlap of all input regions with all intersecting genomic annotations selected by the user, with a user-defined threshold overlap between regions and annotations. The summarize module enables users to quickly compute summarized information of any numerical (Table S2) or categorical data (Table S3) over the annotations or any grouping of categorical data.

The collective goal of the visualization module is to provide insight into modes of regulation, and to discover specific relationships among the input regions and genomic annotations with minimum code or forethought. Consider bisulfite sequencing results from methylSig (Park, 2014) reporting genome-wide differential methylation (DM) between two sample groups. It has columns for DM status, p-value, methylation difference between the groups, and methylation rates of each group. The annotatr package implements functions to show: 1) the number of DM regions in each annotation type with the option to compare against randomized regions (Figure S2), 2) a heatmap of the number of regions annotated to pairs of annotation types (Figure S3), 3) the distribution of numerical data across the annotations or any categorical variable (Figure 1A), 4) the joint distribution of two numerical data columns across the annotations or any categorical variable (Figure S4), 5) the distribution of numerical data for regions in any one two intersecting annotations (Figure 1B), and 6) the distribution of a categorical variable across the annotations or any other categorical variable (Figure 1C).

**Figure 1.**
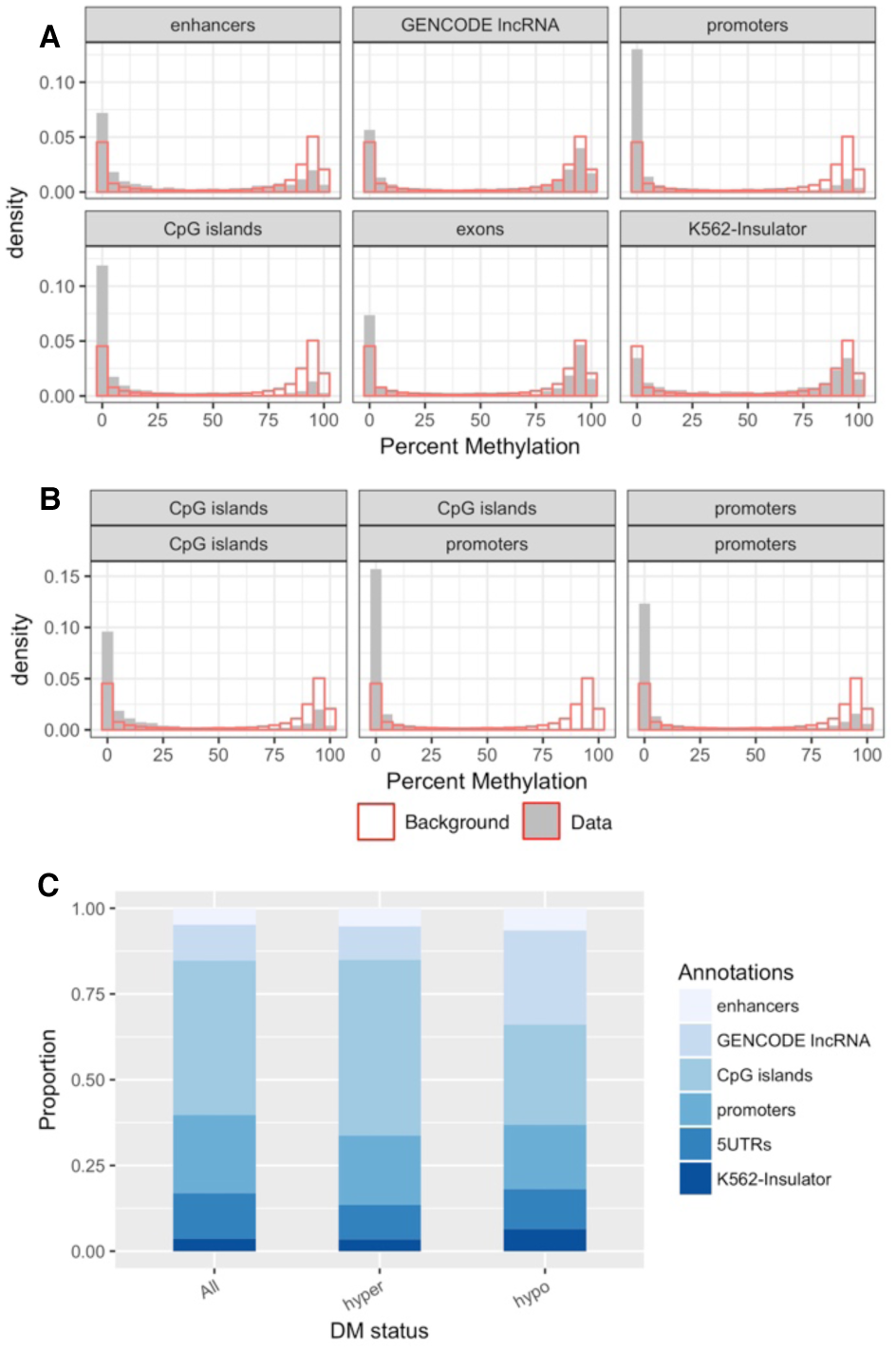
(A) The distribution of the methylation rate across annotations (solid) with the background distribution (outline). Note the clearly visible hyper- and hypo-methylation trends in the different annotation types. (B) The distribution of the methylation rate of regions in just CpG islands (left), promoters and CpG islands (middle), and just promoters (right). Note the relative hypermethylation trend in the co-annotated regions compared to the singly annotated regions. (C) The proportion of annotations of hyper- and hypo-methylated regions, with the background distribution (All) for comparison. Note the differences in enhancers, CpG islands, lncRNAs, and K562-insulators between hyper- and hypomethylated regions compared to each other and all tested regions.

We compared runtimes between ChIPpeakAnno (v3.8.1), goldmine (v1.0.0), and annotatr (v1.0.1) on three data sets varying in size from 31 000 to 2 500 000 lines (Supplementary Methods). annotatr performs up to 13.1x faster than ChIPpeakAnno, and up to 27.5x faster than goldmine, with increasingly better performance as file size increases (Table S4 and Figure S6). In addition to benchmarking, we have compared the features of the three packages (Table S5).

## 3 Discussion

Associating regions of interest to genomic annotations is a standard part of many bioinformatics pipelines. The annotatr package improves upon existing annotation tools by returning *all* the genomic annotations associated with a region instead of artificially prioritizing one annotation type over another, giving a clearer picture of the biological complexities at play. In addition to tabular output of the annotations, annotatr's built-in plotting functions provide an easy and flexible way to summarize the annotations and view how data associated with the regions changes in different genomic contexts. The annotatr package thus enables fast exploration, more complete genomic contextualization of experiments, and more potential discoveries.

## Acknowledgements and Funding

We thank Hani Habra and Jian Kang for their input on the conception and implementation of the package. We also thank Shweta Ramdas and Chee Lee for feedback on the manuscript and package vignette. RC was funded on a Bioinformatics Training Grant (T32 GM070499) and MAS by R01 CA158286 and P30 ES017885.

## Conflict of Interest

none declared.

